# Challenges and emerging directions in single-cell analysis

**DOI:** 10.1101/127761

**Authors:** Guo-Cheng Yuan, Long Cai, Michael Elowitz, Tariq Enver, Guoping Fan, Guoji Guo, Rafael Irizarry, Peter Kharchenko, Junhyong Kim, Stuart Orkin, John Quackenbush, Assieh Saadatpour, Timm Schroeder, Ramesh Shivdasani, Itay Tirosh

## Abstract

Single-cell analysis is a rapidly evolving approach to characterize genome-scale molecular information at the individual cell level. Development of single-cell technologies and computational methods has enabled systematic investigation of cellular heterogeneity in a wide range of tissues and cell populations, yielding fresh insights into the composition, dynamics, and regulatory mechanisms of cell states in development and disease. Despite substantial advances, significant challenges remain in the analysis, integration, and interpretation of single-cell omics data. Here, we discuss the state of the field and recent advances, and look to future opportunities.

## Background

Cell-to-cell variation is a universal property of multi-cellular organisms, which contain diverse cell types characterized by different functions, morphologies, and gene expression profiles. Even within any single tissue, no matter how apparently homogeneous, there is a diverse population of cells, all of which represent different manifestations of that tissue type. Investigation of tissues or cell populations is inherently limited by the fact that the readout of any pooled assay that uses bulk tissue represents a weighted average of that population’s cellular constituents. Intrinsic cellular heterogeneity is obscured in the typical ensemble studies on which the canon of modern biology and medicine are constructed.

Consider, for example, the diverse repertoire of cells present in the three most rapidly self-renewing tissues in mammals: blood, skin, and the intestinal epithelium. Although the trajectory from stem to terminally differentiated cell is almost certainly a continuum of highly variable states, our limited understanding forces us to regard known stem and progenitor cell populations as discrete and stable entities. Even in post-mitotic tissues such as the adult brain, the differentiated cell states resulting from complex bifurcating developmental trajectories may also appear as a continuum. The diversity of cellular states is not only caused by their own inherent cell-to-cell variability, but also influenced by interactions among tens or even hundreds of distinct cells. These considerations question the precise boundary of a cell type and point to the need for single cell analysis to dissect the underlying complexity and the empirical reality of stable and distinct cell states.

The past few years have seen the introduction of technologies that provide genome-scale molecular information at the resolution of single cells, providing unprecedented power for systematic investigation of cellular heterogeneity in DNA [1, 2], RNA[3], proteins [4], and metabolites[5]. These technologies have been applied to identify previously unknown cell types and associated markers [6-8] and to predict developmental trajectories [9-13].

Beyond expanding the catalog of mammalian cell states and identities, single cell analyses have challenged prevailing ideas of cell-fate determination [14-19] and opened new ways of studying the mechanisms associated with disease development and progression. For example, single-cell DNA sequencing (scDNA-seq) has revealed remarkable cellular heterogeneity inside each tumor, significantly revising models of clonal evolution [20-22], whereas single-cell RNA sequencing (scRNA-seq) has shed new lights on the role of tumor microenvironments in disease progression and drug resistance [23].

The ambitious goal of understanding the full complexity of cells in a multi-cellular organism collectively requires not only experimental methods that are considerably better than existing platforms, but synchronous development of computational methods that can be used to derive useful insights from complex and dense data on large numbers of diverse single cells. Several recent papers have discussed various challenges critical to advance the incipient field of single cell analysis [24-27]; here we expand on these discussions with a focus on looking to the future.

## Current challenges in analyzing single-cell data

While many methods have been successfully used for the analysis of genomic data from bulk samples, the relatively small number of sequencing reads, the sparsity of data and cell population heterogeneity present significant analytical challenges in effective data analysis. Recent advances in computational biology have greatly enhanced the quality of data analyses and provided important new biological insights [24-27].

### Data preprocessing

The goal of data preprocessing is to convert the raw measurements to bias-corrected and biologically meaningful signals. Here we focus on scRNA-seq, which has become the primary tool for single cell analysis. Gene expression profiling by scRNA-seq is inherently noisier than bulk RNA-seq, as vast amplification of small amounts of starting material combined with sparse sampling introduce significant distortions. A typical single-cell gene expression matrix contains excessive zero entries. The limited efficiency of RNA capture and conversion rate combined with DNA amplification bias may lead to significant distortion of the gene expression profiles. On one hand, even transcripts that are expressed at a highly level may occasionally evade detection altogether, resulting false-negative errors. On the other hand, transcripts that are expressed at a low level may appear abundant due to amplification biases. These errors artificially inflate the estimate of the cell-to-cell variability. While a number of methods have been developed to address this issue [28-30], managing dropout events continues to be a challenge. Another source of technical variation is the batch effect, which can be introduced when cells from one biological group are cultured, captured, and sequenced separately from cells in a second condition. If a scRNA-seq experiment is designed improperly, the results can be significantly affected by batch effects [31]. Furthermore, high throughput technologies typically involve multiplexing of thousands or more barcode sequences. Error in demultiplexing may be caused by barcode impurities or external background, which has become increasingly challenging as thousands or more cells are multiplexed by recent technologies. Finally, the cell-to-cell variation may also be attributed to cell size, cell cycle state, and other factors that are irrelevant for cell type identification. Statistical models have been developed to remove such confounding factors [27]. Together, these technical artifacts pose important challenges for data calibration and interpretation.

The entanglement of technical and biological variations poses a significant challenge for evaluating data reproducibility. One approach to directly measure technical variability is to use dilute bulk RNA to approximately single cell level (∼10-50 pg of total RNA) [*32, 33*]. However, this approach has at least two significant limitations. First, RNA purification leaves out cellular factors that may impede RNA isolation and amplification. Second, accurate dilution up to single cell levels is technical challenging. Another approach is to use external spike-ins, such as ERCC [34]. However, this approach also has a number of limitations [35]. First, the spike-in probes typically have different molecular properties than the RNA molecules of interest. Second, the spike-in probes interact differently with respect to different molecular biology protocols. Furthermore, the dynamic range of spike-in sets like ERCC is often not optimized for the dynamic range of a typical single cell transcriptome (∼10^3^ to 10^4^). As such, there is a great need to develop better-controlled methods for separating technical and biological variations. Considering these limitations, targeted approaches aimed at precise quantification of key pathways may provide more biological insights in some applications.

### Lack of spatial-temporal context

Single cell DNA and RNA based assays often contain the following steps: cell isolation, cell sorting, and library preparation and sequencing. During this process, cells are isolated from their local environment and destroyed prior to profiling. These “snapshots” lose important contextual information regarding both, the cells’ spatial environment, and the cells’ position within a trajectory of dynamic behavior [25]. Both sources of information are crucial to interpret the precise state of a cell at the time point of its isolation (and usually destruction).

## Future directions

### in situ transcriptomic analysis

To preserve spatial information, transcriptome can be profiled in situ in fixed cells and tissues, using either *in situ* hybridization (ISH) or sequencing. Single molecule florescence *in situ* hybridization (smFISH) provides a powerful tool for detecting individual transcripts [36, 37]. Using super-resolution microscopy [38, 39], this was extended to image over a dozen mRNA *in situ* regardless of transcript density [40]. More recently, a temporal barcoding scheme was developed that scales exponentially with the number of hybridization, called sequential FISH (seqFISH) [41]. In parallel, *in situ* sequencing methods were developed to directly sequence transcripts in tissue sections [42, 43], which has broad coverage but lower efficiency compared to FISH based methods. More recently, a Hamming distance 2 based error correcting barcode system called merFISH [44] was developed and can be applied to long transcripts (>3 kb). This technology has recently been extended to detect 130 mRNA species [45]. Fundamentally, because of high background in tissues, smFISH based methods are difficult to apply directly for detection of mRNAs in tissues.

An amplified version of seqFISH [46], based on hybridization chain reaction (HCR) [47], allows robust detection of mRNAs in tissues and thick cleared brain samples. Combining amplification and a simple one-drop tolerant error correction scheme, this technology was applied to profile up to 249 genes, with each mRNA detected at ∼80% efficiency, in over 15,000 cells in the mouse brain to resolve the structural organization of the hippocampus with single-cell resolution [48]. The authors identified distinct layers in the dentate gyrus corresponding to the granule cell layer and the subgranular zone. They also found that the dorsal CA1 is relatively homogeneous at the single-cell level, while ventral CA1 is highly heterogeneous. For imaging large samples, such as the brain, imaging speed is rate limiting, rather than the switching time between hybridizations. This is because one can toggle between two samples on the microscope, one that is being imaged and another that is being hybridized. Faster imaging modality such as lattice lightsheet [49] and faster cameras can enable higher throughput in the number of imaged cells.

Future work in spatial genomics will take several directions. First, to combine spatial transcriptome data with scRNA-seq data, one can take an approach where cell states are defined by RNA-seq, and then mapped onto the spatial images and transcription profiles determined by spatial transcriptome data [50]. Second, to increase the optical space available in each cell and allow more mRNAs to be resolved spatially, expansion microscopy [51] can physically enlarge the tissue sample. An alternative image correlation approach [52] can also allow dense transcripts to be decoded. Lastly, analysis of *in situ* transcriptomic data requires development of new computational methods, for example, to automatically detect spatial patterns from combination of multiple genes.

### Live imaging transcriptomic analysis

Cellular and molecular behaviors are highly dynamic and constantly changing. These dynamic behaviors greatly complicate the interpretation of snapshot single-cell analyses because individual cells will differ not only in their molecular state from other cells, but even from themselves if analyzed at a different time point [25]. Importantly, these dynamics may not represent noise, but rather a basis for important regulatory mechanisms controlling cell identity, so it is important to quantify dynamic changes and to understand their relevance [53]. Unfortunately, it is also much more difficult than static snapshot analyses. Cells must be kept alive and unchanged during the continuous – and sometimes very long non-invasive analysis of their behaviors. The acquisition, handling and analysis of time-resolved single-cell data then require specialized technical and theoretical approaches. Not only are the requirements for robustness of data acquisition technologies such as live imaging much higher than for snapshot analyses, but the resulting large and complex volumes of data require specialized solutions. These differ from tools available to analyze snapshot data, and often require self-made custom developments. This holds true for the required theoretical algorithms and for user-friendly implementation [25].

### Lineage tracing

The objective of lineage tracing is to label the progeny of individual cells using molecular markers and use such information to reconstruct the developmental trajectories. Recently, high-throughput lineage-tracing methods have been developed by CRSIPR/Cas9-based multiplexing DNA barcodes synthesis [54-58]. These barcodes are stably registered in the genome and inherited during cell division and differentiation. Additional mutations are cumulated in time, through either combinatorial editing at multiple gRNA target loci [54, 55, 57] or by sequential editing at a single locus [56, 58].

In the latter approach, the investigators introduced genetic mutations at the S. *pyogenes* gRNA-encoding sequence to circumvent the requirement of PAM motif in gRNA recognition, enabling the resulting gRNA to repeatedly target its own locus. In addition, the DNA barcodes can be sequenced *in situ*, thereby preserving the spatial information [58]. Some of the aforementioned technologies have been applied to study developmental loci [54, 55, 57] and immune response [56]. In one study [54], the investigators traced the cell lineages in zebrafish, and found that the majority of cells in each organ are derived from a small number of progenitor cells, whereas different progenitors are biased toward different germ layers and organs. Similar results are reported in an independent study [55]. These lineage tracing technologies will likely have wide-range applications in mapping developmental and disease progression trajectories.

### Single-cell multi-omics

While significant effort has been dedicated to improving the quality and throughput of various omic assays, work is also ongoing to develop methods to profile multiple sources of information in the same cells. Multi-omics profiling is valuable for accurate mapping of cell states and can provide insights into the regulatory mechanisms. For example, genomic DNA and mRNA transcripts from the same cells can be quantified by either physical separation [59] or pre-amplification [60], followed by high throughput sequencing. In the former, extracted genomic DNA can be further processed to bisulfite conversion, leading to simultaneous quantification of the methylome and transcriptome [61, 62]. Bioinformatic analysis of the bisulfite sequencing data can further detect genetic information [63, 64]. Protein and transcriptome have also been measured in the same cells [65]. Multi-omic methods applied to single cells have revealed some surprises. For example, profiling DNA and RNA variability in single acute lymphoblastic leukemia cells suggests that genetic heterogeneity is not responsible for the diverse response of drug treatment (Enver, unpublished).

Recent technologies have moved even beyond single cells to investigate sub-cellular localization of biologically active molecules. For example, nanoliter-scale cell fractionation or micro-manipulation has been applied to measure subcellular information within single cells [66]. On a different front, super-resolution imaging has been applied to map the nuclear compartmentalization of chromatin domains [67]. These subcellular data provide new insights into the precise mechanisms of various cellular processes. Ultimately, we may be able to understand phenotypic differences between genetically identical cells in terms of such variations in subcellular organization.

### Modeling and predictions

Different cell types usually arise from a linear hierarchy of differentiation stages, and one goal of single-cell analysis is to identify previously unknown cell types and lineage relationships. Numerous methods have been developed to isolate similar cell types from single-cell gene expression data [7, 50, 68, 69]. Furthermore, additional methods have been developed to specifically detect rare cell subpopulations [70, 71]. To compensate the dropout effect, methods have also been developed to impute gene expression based on similar cell types [72].

Single-cell analysis has helped refine traditional views of cell differentiation. For example, A number of studies [14-16, 73] report evidence to suggest that megakaryocytes emerge at a “high” level, approximating that of the hematopoietic stem cell (HSC); this insight challenges the prevailing model that the megakaryocytic lineage emerges late in the differentiation cascade. The cell states defined by transcriptomic patterns are surprisingly continuous instead of forming distinct, transcriptionally defined groups [15, 74]. This apparent continuity of cell states poses practical challenges for cell annotation, and conceptually, implies a need for significant revisions to current models of cell lineage hierarchy.

Data from single-cell studies have enabled the development of mathematical models that represent the distribution of cell states as one sampled from a dynamical system [11, 17, 19, 73]. In this view, cell types are modeled as “attractors” [75], stable states that are determined by the underlying gene regulatory networks and sometimes referred to as the energy landscape. In some models, stochastic fluctuation, either due to intrinsic or extrinsic noise, may facilitate dispersion and transition between attractors [11]. Although complex, these mathematical models can be used not only to explain the continuity of cell states but, in some cases, predict the initiation events during cell differentiation, thereby providing mechanistic insights [11]. In a similar way, the hierarchy of cell states can be measured by entropy, which has been applied to inform cell differentiation directions [76, 77]. These new methods have opened up new ways to think about cell states, not as discrete entities, but as a continuum. To connect these two viewpoints, it is critical to determine with high precision the level of natural variation that defines the same cell type and distinguish this from the changes linked to functional state transitions. A major obstacle for achieving this goal is that the resolution of cell-state identification is limited by the quality of the underlying scRNA-seq data, which varies greatly depending on sequencing depth and other factors. Such differences have contributed to the debate over the organizing structure of hematopoietic lineage hierarchy [15, 16].

### Functional validation

As single cell data continue to grow in quality and quantity, new cell states, lineages, and associated markers are being identified at an accelerated rate. It is important to recognize that such findings are typically based on correlative analyses and that their functional relevance needs to be carefully evaluated through further experimental validation.

A first level of validation is to utilize the identified markers to label the predicted cell-type and visualize in their original tissue. For example, unsupervised clustering of 25,000 single-cell transcriptomes identified 15 types of bipolar neurons [8]. The authors identified cell-type specific markers and fluorescently labeled the predicted cell-types by DNA FISH. They found that the spatial organization of these predictive cell-types is restricted to definitive layers and that different cell-types display distinct morphology, thereby supporting their functional identity.

A deeper level of validation requires design of functional assays to demonstrate that a predicted cell type has unique properties. For example, single-cell analysis showed that common myeloid progenitors (CMP) occur in two varieties associated with differential expression of CD55 [14]. Using an *in vitro* colony forming assay, the investigators found that CD55+ CMP produce predominantly erythroid and megakaryocytic (MegE) colonies, whereas few MegE colonies are formed from CD55- CMP, indicating these two subpopulations are functionally different. A similar strategy has been applied to compare the functional difference between HSC subpopulations, termed MolO and NoMO respectively [78]. These investigators found that MolO cells were enriched for higher than average CD150 and Sca-1 surface marker expression and lower than average CD48 expression.

In the same vein, engineered animal models can allow isolation of cell populations and functional testing. For example, comparative scRNA-seq analysis between HSCs from young and old mice identified a gene signature associated with the MegE lineage [79]. By using a transgenic mouse strain carrying a VWF-EGFP reporter, the authors verified an increased bias toward platelet-priming HSCs in old mice.

### Combining scRNA-seq and CRISPR/cas9 based perturbations

CRISPR/Cas9 based genetic screens have been widely used to systematically characterize gene functions [80, 81]. Recently, this technique has been combined with scRNA-seq analysis [82-85], thereby greatly increasing the throughput of functional readouts. In these studies, gene activities are disrupted by either genetic mutations [82, 83, 85] or epigenetic inhibition [84]. gRNA-specific reporter transcripts are synthesized, which can be detected along with the mRNAs by scRNA-seq sequencing. By varying the concentration of the gRNA-containing vectors, the technique can be used to study the gene function either in isolation or in combination. In one study [82], the investigators applied this technology to analyze the effects of 24 TFs in mediating the immune response of dendritic cells. They found that the TFs form distinct modules each targeting a common set of gene program. Further analysis detected significant genetic interactions among a subset of TFs. In another study [83], the engineered hematopoietic progenitor cells were injected into wild-type recipient mice to evaluate their effect in hematopoiesis. This allowed them to identify a previously unknown role of Cebpb in regulating the balance between dendritic cells and monocytes during development. The combination of genome editing and scRNA-seq profiling provides a powerful tool for high-throughput dissection of gene functions and will have a wide range of applications in biomedical research.

### Disease applications

Genomic profiling has been widely used to identify markers, mechanisms, and therapeutic targets of diseases. Most studies to date identify disease related alterations by comparing genomic profiles obtained from bulk disease samples and their normal counterparts. However, these average profiles provide a distorted view of the disease sample if it contains significant cellular heterogeneity, as in cancer. Single-cell technologies have provided a set of powerful tools to dissect the cellular heterogeneity and led to important discoveries in cancer [23, 86-89] and other diseases [90, 91]. For example, multiplexing qPCR analysis has identified subtypes of leukemia cells with distinct capacity of proliferation [87, 92]. Application of scRNA-seq to cancer also led to identification of rare subpopulations associated with drug resistance [23] or self-renewal [89], whereas scDNA-seq can be used to reconstruct paths of clonal evolution [21]. Single cell profiling will provide new opportunities for mechanistic understanding of the initiation and progression of human diseases and to develop novel treatment methods targeting specific cell types.

### Interdisciplinary research

We recognize that to overcome each challenge requires significant resources of lab and computational infrastructure. To move forward, the field needs groups of people with diverse expertise to work together. Interdisciplinary approaches are recognized to be important, if not a crucial prerequisite, for addressing many open questions, but also come with numerous challenges. First, the lack of expertise for parts of an interdisciplinary collaboration requires increased effort and tie for communication. The importance of a common language is well known, but remains a significant problem in almost every new project. Ideally, this hurdle will be overcome by a new generation of students and postdocs who are educated in multiple disciplines like biology/medicine, engineering and theoretical sciences. Interdisciplinary science, while leading to higher long-term impact, tends to be slower and published in journal of lesser impact [93], and is hard to organize and fund. Thus, it needs more patience, in particular in environments with funding cycles that require fast short-term output. Finally, not only language, but also career paths, scientific and publication cultures, hiring procedures and age, and scientific talent and academic motivation vary widely across disciplines. While many of these differences pose managerial challenges and should not impact scientific merit, in reality they often are the reason for failures of interdisciplinary endeavors. Overcoming these problems will require changes in teaching, funding, publication and hiring procedures, which would benefit most areas of science, but will only have a measurable effect after a few years.

## Conclusions

Single-cell analysis is an exciting and rapidly expanding field that holds tremendous potential to improve our understanding of fundamental biological problems and to better understand the nature and complexity of human disease in order to develop more effective therapies. To achieve these ambitious goals, proper control needs to be taken to warrant the detection of genuine heterogeneity existing in cell population and tissue samples. In addition, we need to invest in development of new methods. Single-cell data presents a number of intrinsic challenges, including systematic noise, the features of biological systems, and the sparsity and complexity of the data. The past few years have witnessed remarkable growth in the field, a trend we believe will continue, enabling more rigorous development of methods and deeper understanding of biological complexity.

## Acknowledgement

This article is a result of the discussions at the Radcliffe Institute Exploratory Seminar on “Theoretical Challenges in Single-Cell Analysis” in June 2016. We are grateful for the Radcliffe Institute’s generous financial and logistical support.

## Authors’ Contributions

GCY conceived the study. All authors participated in the discussions and writing of the manuscript.

